# Context- and sex-dependent links between sire sexual success and offspring pathogen resistance

**DOI:** 10.1101/2024.09.10.611971

**Authors:** Aijuan Liao, Tadeusz J. Kawecki

## Abstract

Sexual selection has been proposed to promote genetic variants that improve resistance to pathogens (a special case of the “good genes” hypothesis). Yet, experimental tests of this hypothesis are scarce and equivocal. It is often assumed that additive genetic correlation between sexual traits and pathogen resistance is generated by their shared dependence on genetically variable “condition” of the organism. However, an alternative scenario posits condition-independent genetic variation in pathogen resistance; individuals more resistant to currently prevalent pathogens remain healthier and can invest more in sexual traits, but this advantage disappears in the absence of pathogens. Here, we tested whether *Drosophila melanogaster* males that are more sexually competitive (in terms of paternity share) sire offspring that are more resistant to the fungal pathogen *Metarhizium brunneum*. Furthermore, to investigate the importance of epidemiological context, we exposed sires to either an infection or a sham treatment before mating, to test whether the sire-offspring relationship depends on the presence of pathogens during sexual selection. We found that the relationship between sires’ sexual success and offspring pathogen resistance not only depended on sires’ exposure to the pathogen, but also on offspring sex. Sires that were more sexually successful in the absence of the pathogen had less resistant offspring whereas no relationship was detected for sires that competed for paternity after pathogen exposure. For daughters, the relationship tended to be negative irrespective of sire’s pathogen exposure. In no case was a positive correlation predicted by the “good genes” hypothesis detected. Thus, while sexual selection may act on genes affecting resistance in a context- and sex-dependent manner, we found no circumstances under which it promoted resistance.

**Lay Summary:** Sexual selection has been debated in relation to natural selection for decades. The “good genes” hypothesis which posits a positive genetic correlation between sexual success and non-sexual fitness, is often considered as a mechanism aligning sexual selection and natural selection. Scientists have long wondered if genetic variants that make individuals more sexually successful also enhance their ability to fight against pathogens, testing a special version of “good genes”. However, evidence is mixed. Two key mechanisms linking sexual success and pathogen resistance have been proposed: the “condition- dependent” scenario, where general health improve both sexual traits and pathogen resistance, and the “context-dependent” scenario, where resistance to specific pathogens benefits sexual success only in certain environments. Few studies distinguish between these two mechanisms. This study examines both scenarios in *Drosophila melanogaster*, finding that the relationship between a sire’s mating success and offspring pathogen resistance to the fungal pathogen *Metarhizium brunneum* depends on sire’s pathogen exposure prior to measurement of sexual success and offspring sex. These findings challenge the "good genes" hypothesis and emphasize the necessity of including context in sexual selection studies.

## Introduction

Sexual selection, initially proposed by Darwin (Darwin, 1871), has been a topic of ongoing debate over the years, especially its relationship with natural selection. The triumph of the sexiest is often observed going against the survival of the fittest. For example, calling in frogs decreases their ability to avoid predators, elongated tails in birds reduce their foraging rates, and so on (Andersson, 1994). However, sexual selection can also reinforce natural selection (Kokko et al., 2002; Hollis et al., 2009; Parrott et al., 2019) if there is a positive additive genetic covariance between sexual success and non-sexual fitness traits. This is the essence of the “good genes” hypothesis (Zahavi, 1975; Møller & Alatalo, 1999; Houle & Kondrashov, 2002). It suggests that alleles favored by sexual selection also improve offspring survival or other non-sexual aspects of offspring fitness. One such non-sexual fitness trait often of interest in the context of natural and sexual selection is pathogen resistance, the ability of an individual to fight against or tolerate the negative impacts of pathogens. Existing empirical data provides equivocal evidence for the genetic correlation between sexual success and pathogen resistance (Folstad & Karter, 1992; Kruuk et al., 2002; Hall et al., 2004; Prokop et al., 2012; Wu et al., 2018; Parrett et al., 2022), indicating a complex relationship between these fitness-relevant traits.

Two mechanisms have been proposed on how a genetic correlation between pathogen resistance and traits that affect sexual success can be generated; they differ in the assumptions on the genetic architecture of resistance and the importance of the environment context (summarized in (Westneat & Birkhead, 1998)). The first proposed mechanism posits that sexual selection favors general immunocompetence which is effective against a broad range of pathogens. Within this theoretical framework, the level of both pathogen resistance and sexual trait expression are largely determined by the individual’s condition (hereafter, “condition-dependent” scenario). Here, condition refers to the general physiological state of the individual and the ability to maintain homeostasis in face of environmental challenges (Hill, 2011), a latent trait hypothesized to show appreciable genetic variation because it captures much of the mutational variance across the genome (Rowe & Houle, 1996). The common condition-dependency generates a positive genetic correlation between sexually selected traits and resistance to a range of pathogens. In this scenario, the direction of the relationship between sire sexual success and offspring pathogen resistance is not affected by the epidemiological context. As a consequence, sexual selection would be generally expected to promote resistance to a variety of pathogens.

Under the second scenario, genetic variants confer resistance to specific pathogens largely independent of the individual’s general condition (Hamilton & Zuk, 1982). Instead, condition is assumed to be strongly affected by the degree to which an individual can avoid or resist currently prevalent pathogens. Hence, males carrying genotypes conferring resistance to currently prevalent pathogens will be in better condition, be able to invest more in sexually selected traits and enjoy a higher sexual success than males endowed with less resistant genotypes. However, if the pathogen is absent, genes for resistance to it bring no benefits and may even inflict some physiological cost due to investment in immune system or collateral damage caused by it), resulting in lower expression of sexually selected traits compared to males carrying susceptible genetic variants (Adamo & Spiteri, 2005). Thus, the sign and the strength of genetic correlation between pathogen resistance and sexual traits is mediated by prevalence of pathogens (i.e., the epidemiological context), a form of genotype by environment interaction (hereafter, “context-dependent” scenario). As a consequence, sexual selection would only be expected to promote resistance to currently prevalent pathogen(s) and would possibly counter-select against resistance in their absence.

Nearly all of the rather few experimental studies explicitly testing the relationship between male’s sexual traits or sexual success and offspring pathogen resistance implicitly or explicitly assumed the “condition-dependent” scenario, and thus did not expose the males themselves to the pathogen (Achorn & Rosenthal, 2020). To our knowledge, only one study was explicitly designed to distinguish between these two scenarios Joye and Kawecki (2019). Using *Drosophila melanogaster* and the bacterial pathogen *Pseudomonas entomophila*, they showed that males winning a mating contest sired more pathogen-resistant sons than loser males if they had been exposed to pathogens prior to mating contest. However, the reverse was true when the pathogen was not present, supporting the latter “context-dependent” scenario. A limitation of that study is that the winner-loser metric only scored who mates first with a single virgin female, which may not represent the overall net results of sexual selection. The genetic correlation between sexual success and non-sexual fitness varies across different aspects of sexual selection (Rowe & Rundle, 2021). The selection for good genes can be rather stronger if multiple episodes of sexual selection (e.g., mating success, fertility, mate monopolization, sperm defense or offense) act on promoting good genes (House et al., 2016; Vuarin et al., 2019) or of little overall net effect if the cost of sex-specific selection or sexual conflict outweighs the benefits (Pizzari & Birkhead, 2000; Okada et al., 2014; Baur et al., 2024).

In this study, we aimed to assess the genetic relationship between offspring pathogen resistance and a measure of the sire’s sexual success – competitive paternity share – that reflects the net effect of the diverse facets of sexual selection. By exposing the sires either to the pathogen or to a sham treatment, we tested whether this relationship depends on the epidemiological context under which sexual selection occurs, as proposed by the “context-dependent” scenario described above. Our experiment system consisted of the fruit fly *Drosophila melanogaster* and *Metarhizium brunneum*, a fungal pathogen that develops relatively slowly relative to the life cycle of *Drosophila* (median time to death 7-9 days), even though the immune system is already strongly activated after 2 days (Liao et al., 2024). This experimental system allowed us to compare a situation where the pathogen burden only becomes significant several days after the mating group has been put together (“long mating” scenario) with the case where the disease is already advanced by the time naïve sires first encounter virgin females (“short mating” scenario). These two mating scenarios (described in detail below) are expected to differ with respect to relative importance of mating success versus sperm competition and its prevention (Rice & Chippindale, 2001; Pischedda & Chippindale, 2006).

In addition to testing for the effect of epidemiological context, we also tested for differences in the relationship of sire’s sexual success with pathogen resistance of sons versus daughters. Pathogen resistance often differs between sexes (Belmonte et al., 2019), and the fact that sex affects the expression of many genes (Ellegren & Parsch, 2007; Ayroles et al., 2009; Innocenti & Morrow, 2010; Ingleby et al., 2014; Duneau et al., 2017) suggests the possibility of genotype by sex interaction for pathogen resistance (McKean & Lazzaro, 2011; Klein & Flanagan, 2016). If so, genetic variants favored by sexual selection may have sex-specific or sexually antagonistic effects on pathogen resistance. Indeed, a previous study found a relationship between sire’s winning a mating contest and offspring resistance for sons, but not for daughters (Joye & Kawecki, 2019). Thus, what is a “good gene” may depend on the sex, further complicating the predictions about the effect of sexual selection on non-sexual traits.

## Materials and Methods

### Fly origin and maintenance

Wild-type flies used in the experiment originated from a lab-adapted, outbred population collected in 2007 in the Canton of Valais, Switzerland (Valais 07) and maintained in the lab since at a population size of more than 1000 adults with overlapping generations. To generate a fluorescently labeled competitor strain, we backcrossed GFP-ProtB (courtesy of Stefan Lüpold, University of Zürich, Zürich, Switzerland (Manier et al., 2010)) into this background (Valais 07) for five generations. GFP-tagged flies used in the experiment were more than 95% genetically identical to the wild-type population. The ubiquitously expressed GFP is a very effective dominant phenotypic marker for paternity assignment as it is already visible at the embryo stage.

Adult flies used in this experiment were raised on standard cornmeal-yeast-agar media with Nipagin at 25°C, 55% relative humidity, and 12L:12D photoperiod. Nipagin (Methylparaben) is an antimicrobial agent that prevents fungal growth on the fly food medium, but does not prevent *M. brunneum* from proliferating inside fly bodies. Larval density was controlled by transferring about 200 eggs to each bottle containing 40ml food. Virgin wild-type flies of both sexes were collected within 6-8h post emergence and then maintained in same-sex groups of 10 in the food vials with 10ml food until used in the experiment. GFP-tagged non-virgin males were collected three days after emergence and also maintained in groups of 10. All fly transfers were done under light CO_2_ anesthesia.

### Fungal culture and infection protocol

The pathogen used in this experiment is *Metarhizium brunneum* KVL 03-143. Pathogen origin and infection protocol were previously described in Liao et al. (2024). Sires assigned to the infection treatment were individually dipped into 2ml spore suspension (spores stored in 0.05% Triton X-100) with an adjusted concentration of 10^7^ spores/ml for 10 seconds. Sires assigned to sham treatment were individually dipped into spore-free 0.05% Triton X-100 for 10 seconds. To study the offspring resistance, offspring of the same family were submersed in the spore suspensions (10^7^ spores/ml) in groups of 20 for an extended 30 seconds to make sure that all individuals were exposed to the spores. Infection treatment was applied to daughters and sons separately.

### Experiment design

The design of our main experiment is summarized in Figure 1.

**Figure 1.**
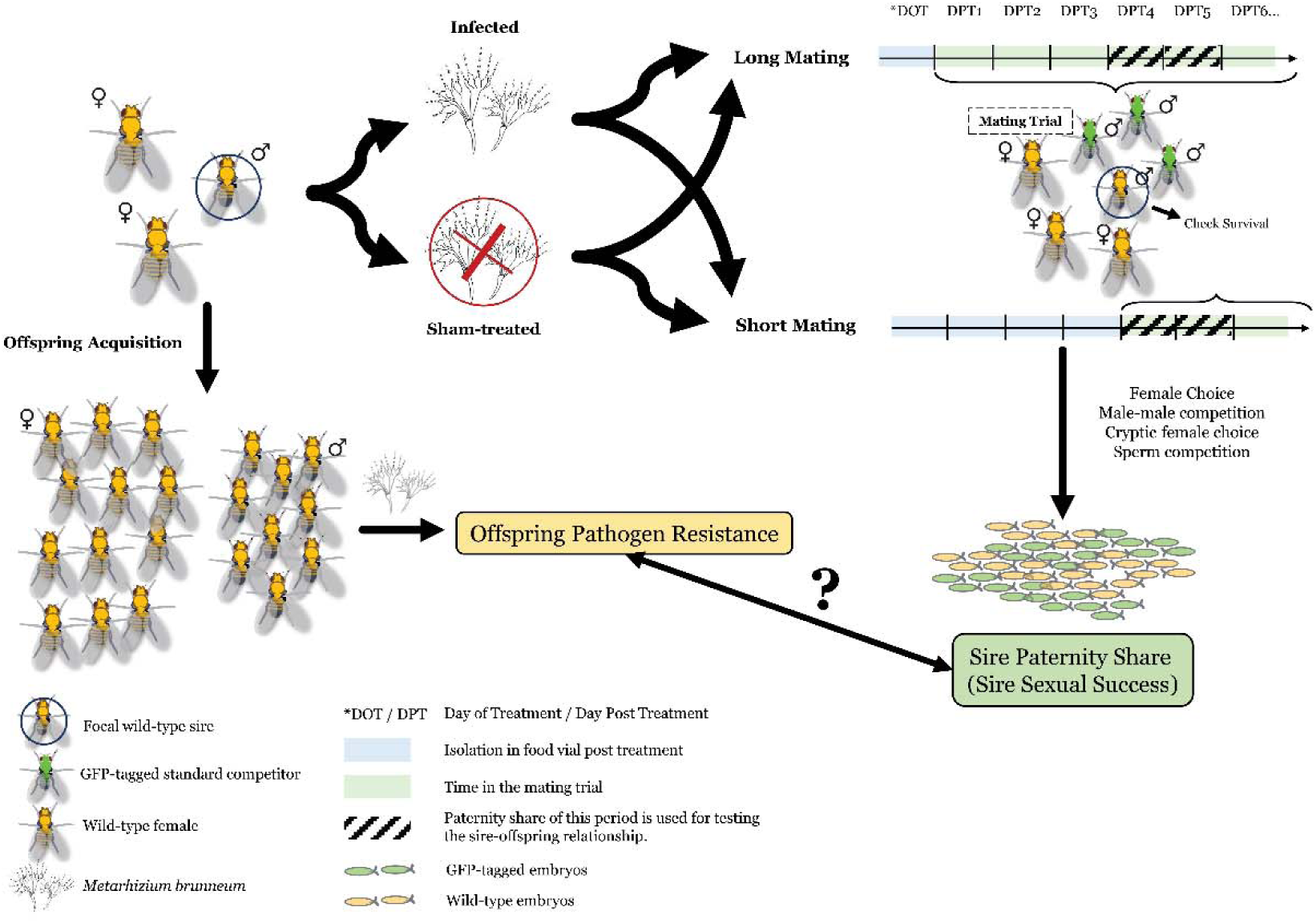
Experimental design to study the relationship between sire sexual success and offspring resistance to Metarhizium brunneum with/without exposure to the pathogen before sire sexual success is assessed.

Our aim was to investigate the relationship between a measure of sexual success for each sire and a measure of pathogen resistance of each sire’s offspring of either sex, depending on the prior exposure of the sires to infection. To rule out any potential effect of sire treatment on the measurement of offspring’s resistance, we obtained offspring from each sire prior to any treatment. To this end, each two-day-old virgin male (sire) was paired with two random wild-type virgin females in a vial with standard food and given 24 hours to mate before being removed for the next step of the experiment. Females were given another 48 hours to lay eggs in the same vial before being removed. All vials at this step (offspring acquisition vials) were kept for offspring collection.

Subsequently, we quantified the sires’ sexual success following either infection or sham treatment. Six hours after being removed from the offspring acquisition vials, each sire was subjected at random to either infection or sham treatment and then kept individually in a food vial for either 1 or 4 days (see below) until the mating trial. In each mating trial, a single sire was placed in a bottle with grape juice agar and yeast (i.e., oviposition medium) together with three virgin females and three GFP-tagged competitor males (neither subject to infection). GFP-tagged males outnumbered focal males in the mating trial because they are less competitive in sexual competition with wild-type males (Sharda et al., 2024). Furthermore, using multiple competitors and mates was expected to reduce the impact of individual variation among them on the measure of sexual success of the focal sire. Upon the onset of the mating trial, the virgin females were 4 days old; for workload reasons, the competitor males were not isolated as virgins and were of the same age as the sire. Flies in the mating trial were transferred to a new oviposition medium following a 10h (daytime):14h (nighttime) interval. An extended nighttime interval allows us to cover both the morning and evening peaks of egg laying. All embryos laid during the nighttime hours were counted (by a single experimenter blind to sample identity) under fluorescence stereomicroscope to acquire the number of GFP-tagged and wild-type embryos (the percentage of non-GFP embryos used as the paternity share of the wild-type sire).

To study if the temporal dynamics of interactions within the mating trials and the sire’s infection status upon the onset of the mating trial affected the relationship between the sire’s sexual success and offspring pathogen resistance, we designed two mating scenarios: “Long Mating” and “Short Mating”. Sires assigned to the “Long Mating” were placed with virgin females and competitor males 1 day after the infection (or sham) treatment; thus, by the time the infection developed all females would have mated multiple times. In the "Short Mating", the sires were only placed with virgin females and competing males 4 days post infection/sham treatment. In both cases, we used the paternity of offspring produced by the females on days 4-5 after the infection or sham treatment of the focal sires. This allowed the infection to develop and the immune system to become fully activated (Liao et al., 2024) but minimized losses due to sire mortality. Thus, in the "Short Mating" treatment the females were virgin and males sexually naïve at the beginning of the period of scoring the paternity, whereas in the "Long Mating" scenario the females were likely multiply mated and having experienced three days of sexual and competitive interactions. In the latter case, some offspring produced during the focal period may have resulted from mating that occurred earlier. On day 7 post-treatment, all flies in the mating trials were once more transferred to standard food bottles, and their mortality was recorded daily until day 10 post treatment. No mortality of the sham-treated sire nor fungal infection of the females in contact with infected males was observed during the experiment.

To assess the offspring pathogen resistance, from each sire, we randomly collected 20 4-6-day-old non-virgin offspring of each sex and infected them with *M. brunneum* spore suspension (10^7^ spores/ml). After infection, offspring were kept in same-sex groups of 10 in the food vials. Mortality due to infection was then recorded daily until day 9 post infection. Flies found dead within 2 hours post infection were considered as mortality from handling rather than infection and were removed from the vials and the analysis, leaving 7551 offspring of 111 infected and 101 sham-treated sires for the final analysis). The entire experiment was conducted in three blocks over two months (see Supplementary Table S1 for detailed experiment setup).

We chose paternity share of day 4-5 post treatment as a proxy for sire sexual success because at this time the infection is established, the immune response is fully activated, and mortality of infected males usually starts on day 6 post infection (Liao et al., 2024). The paternity share was calculated as the sum of the wild-type egg count from day 4 and day 5 divided by the total number of eggs (wild type + GFP-tagged) laid in that period. Sires that died before day 6 post infection were removed from the final analysis.

To further investigate sire’s overall sexual competitiveness and to be able to attribute any observed relationships to sexual selection, within the main experiment work, we also recorded the paternity share of sires in the “Long Mating” scenario from day 1-6 post treatment and that of sires in “Short Mating” scenario from day 4-6 post treatment. Sires found dead during the mating trials were removed from the arena and we have stopped tracking the paternity share of the corresponding sire since the day of mortality.

### Statistical analysis

All statistical analyses were done with R (v. 4.1.2) (R Core Team, 2020). Visualization of the results was conducted with package *ggplot2* (v.3.4.1) (Wickham, 2016) and *ggpubr* (v.0.4.0) (Kassambara, 2020). All generalized linear mixed models (GLMM) were performed with package lme4 (v.1.1-27.1) (Bates et al., 2015), and the continuous covariates in the model were mean-centered (i.e., subtracting the overall mean value across all experiment blocks). We used the *DHARMa* package (v.0.4.5) (Hartig, 2022) to check for distribution of the residuals and overdispersion. Effects of the fixed factors in the GLMMs were subject to likelihood ratio test (LRT) with the *mixed()* function of the *afex* package (v.1.0-1) (Singman et al., 2021). Major axis regression was performed with *lmodel2* package (v.1.7-3) (Legendre, 2018).

#### Effects of the pathogen on sire’s sexual success in a competing environment

We compared the paternity share (the number of offspring sired by the focal sire versus that by the standard competitors) in the mating trial of the infected sires to that of the sham-treated sires. We fitted the data to a GLMM (binomial distribution, logit link) with day post treatment (a continuous variable), sire infection treatment (infected vs. sham-treated), their interaction and experiment block (N = 3) as fixed factors, and sire identity as the random factor. The analysis was done for data from days 4-6 post treatment and for the "Long Mating" and "Short Mating" scenarios separately.

To obtain unbiased mean estimates of paternity share of the sires, we fitted, separately for each sire infection treatment and day post treatment, a random-effect-only binomial model with logit link, using the number of offspring sired by the focal sire versus that by the standard competitors as the response variable, and sire identity and experiment block as random factors. Back-transformed (inverse logit) intercept and its standard error of these models were used as estimates of expected paternity (mainly for plotting purposes). An analogous approach was used to estimate the mean survival of offspring to a particular day post-infection.

#### Offspring survival post infection

To test if there is any difference between the post infection survival of sons and daughters, we performed a GLMM (binomial distribution, logit link) with the number of alive offspring versus the number of dead offspring as the response variable, day post treatment (a continuous variable), offspring sex and their interactions, and experiment block as fixed factors and sire identity as the random factor.

In addition, we fitted linear models to investigate the relationship between the daughter’s and son’s survival on each day post infection, and the correlation was tested with Pearson’s r correlation method. To include the block effects in the survival data, the offspring survival was centered on zero by experiment block before feeding into the linear models.

While the assay of offspring survival post-infection yielded survival curves, the analysis of the sire-offspring relationship required a single measure of pathogen resistance per family per sex. As such a measure, we used the offspring survival data (the number of alive flies versus dead flies) from the day when the average proportion of alive offspring across all families dropped down to around 50%, and the chosen day could be different for daughters and sons. To examine the relationship between daughter’s pathogen resistance and son’s pathogen resistance, we performed a major axis regression (MA) (Warton et al., 2006) based on the assumption that the strength of prediction is the same when using daughter pathogen resistance to predict son pathogen resistance and when vice versa. The regression line was not forced through the origin to capture the sex differences in the susceptibility to the pathogen. In addition, we tested the correlation with Pearson’s r correlation method.

#### Sire’s sexual success, sire’s pathogen resistance and offspring’s pathogen resistance

Before examining the relationship between sire sexual success and offspring pathogen resistance, we first investigated whether there is a link between sire sexual success and sire pathogen resistance and investigated the heritable potential of pathogen resistance.

We first examined the relationship between sire’s sexual success and sire’s pathogen resistance. For this analysis we focused on sires that were subject to the infection treatment, which we divided into two classes, relatively resistant (i.e., those alive after the day when 50% mortality was observed) and relatively susceptible (i.e., those that died before or on the day when 50% mortality was observed). Sires that died prematurely (i.e., before day 6 post infection) were excluded from the analysis. We did not use the actual day of death because ∼25% of sires survived until day 10 (the last day of the experiment) and would thus have been censored. The splitting time points for the two mating scenarios are day 9 post infection for the “Long Mating” scenario and day 7 post infection for “Short Mating” scenario (Supplementary Figure S1). We fitted the data with a GLMM (binomial distribution, logit link), including sire paternity share as the response variable, sire pathogen resistance level (relatively resistant versus relatively susceptible) and experiment block as fixed factors, and observation identity as the random factor to correct for overdispersion.

We also investigated the relationship between the sire’s pathogen resistance level and the offspring’s pathogen resistance. While this should reflect the heritability of pathogen resistance, it should be noted that the trait was measured in sires and offspring under different circumstances and on a different scale. We fitted the data to a GLMM (binomial distribution, logit link), using offspring pathogen resistance (number of alive offspring versus the number of dead offspring on the chosen day) as the response variable, sire pathogen resistance level and experiment block as fixed factors, and observation identity as the random factor to correct for overdispersion. The analysis was done for daughters and sons separately.

We applied the same set of analyses to the dataset where sires were divided into six levels based on their mortality time. Sire pathogen resistance level was included in the GLMM either as a continuous variable or a categorical variable, and similar conclusions were reached (results not shown).

#### Finding “Good Genes”: Relationship between sire’s sexual success and offspring’s pathogen resistance

To address the central question of this study – the relationship between sire sexual success and offspring pathogen resistance – we fitted the data to a GLMM with binomial distribution and a logit link function. The full model included offspring pathogen resistance as the response variable, sire paternity share, sire infection treatment, mating scenario, offspring sex and their interactions, and experiment block as fixed factors, sire identity as the random effect, and an observational level random factor (i.e., offspring vial ID) to correct for overdispersion.

Based on the interactions observed in the full model, we split the analysis by offspring sex, resulting in GLMMs with sire paternity share, sire infection treatment, mating scenario and all the interactions, and experiment block as fixed factors and sire identity as the random factor. We then performed model selection and chose the model with the lowest AIC as the best model describing the data. Estimates of the relationship (i.e., slope) between offspring’s pathogen resistance and sire’s sexual success and the associated standard errors were from GLMMs with sire paternity share and experiment block as the fixed factor, and sire identity as the random factor.

## Results

### Effects of the pathogen on sire’s sexual success in a competitive environment

Paternity share in a competitive environment serves as a proxy for the focal sire’s sexual competitiveness. In the mating trial, each focal sire was outnumbered by the GFP-tagged males (1 vs. 3), but on average half of the embryos were sired by the focal males, which confirms that GFP-tagged males are inferior as sexual competitors, reflecting a cost of the GFP construct (Sharda et al., 2024). The paternity share was highly variable among individuals, ranging from zero to 100% for both infected and sham-treated sires. In the Long Mating scenario, where the mating trial started on day 1 after infection or sham-treatment of the focal sires, the dynamics of paternity share on days 1-3 in the mating trial (i.e., days 1-3 post infection) was different from that on days 4-6 (Figure 2A). On days 4-6 the paternity share of both infected and sham-treated sires decreased, with infected sires experiencing a greater decrease (LRT, sire infection treatment × day post treatment, χ^2^_1_ = 43.5, *p* <0.001; Supplementary Figure S2; Supplementary Tables S2, S3). In the Short Mating scenario, the mating competition started on day 4 post-treatment, and infected sires entered the competition when the pathogen was already well established and the immune system was strongly activated (Liao et al., 2024). Since the start of the mating trial, infected sires had a lower paternity share than the sham-treated sires. As in the Long Mating scenario, paternity share decreased over time, and different rates of decrease were detected for infected and sham-treated sires (Figure 2B; sire infection treatment × day post treatment, χ^2^_1_ = 12.3, *p <* 0.001; Supplementary Tables S2, 3). These findings indicate that infected males could still court and mate, but infection negatively affected their competitiveness. Paternity share scored on a given day does not necessarily reflect the mating on that day because females can store sperms and become less receptive once mated. In the subsequent analyses, we used paternity share on day 4-5 post treatment as the measure of the sire’s sexual success – it captured the treatment effects in both mating scenarios and no major sire mortality was observed during this timeframe.

**Figure 2.**
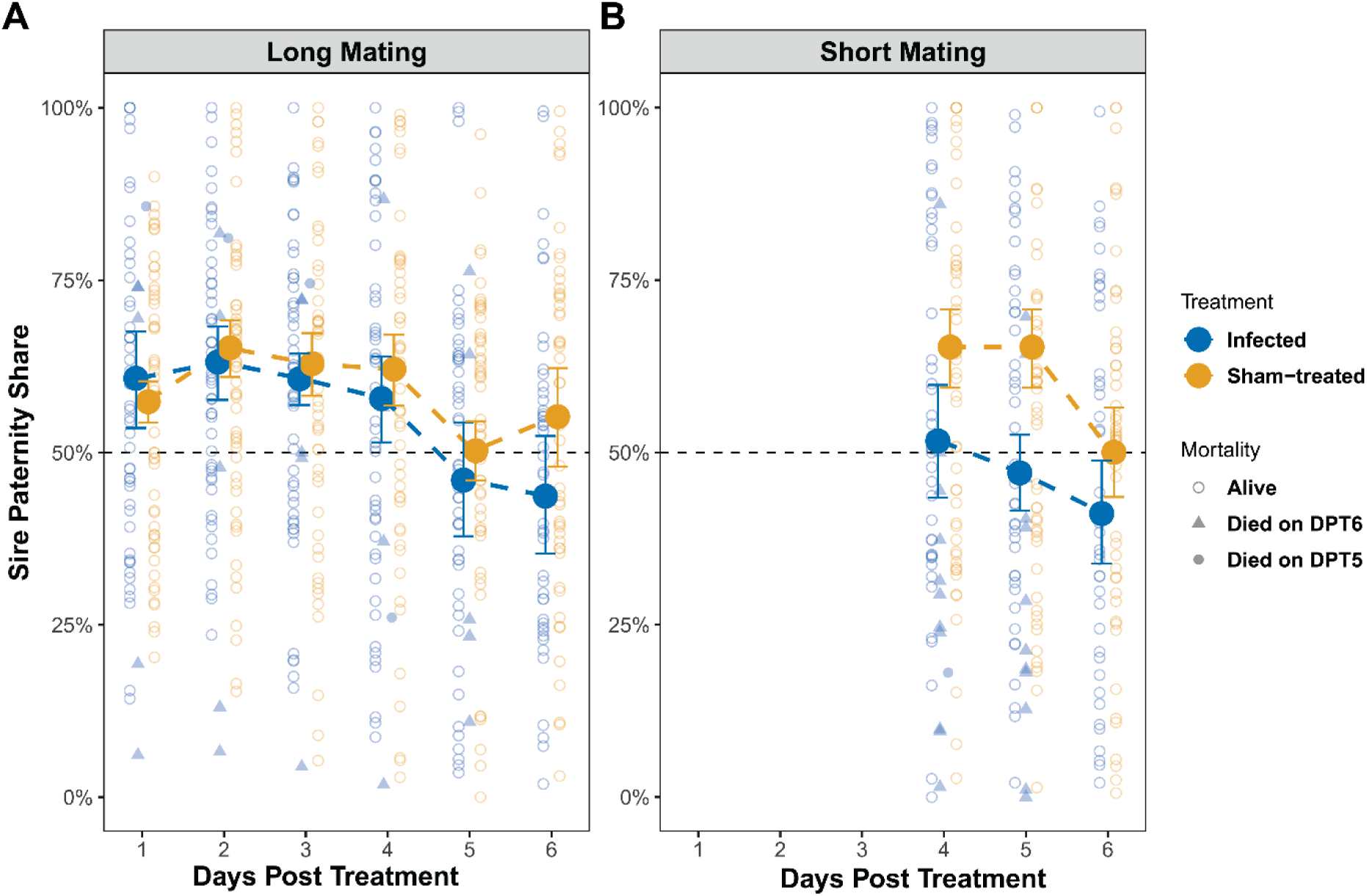
Paternity share of the infected sires and sham-treated sires in A:”Long Mating” mating trials and B: “Short Mating” mating trial. Large symbols represent estimated means ± SE. Estimated means are extracted from random-effect-only GLMMs (see Methods). Small symbols are values for individual sires, symbol shape indicates on which day post treatment the sire was found dead (“Alive” means alive until the end of day 6). Paternity share for a given day is only plotted if the sire was alive at the end of the day. One infected sire that died before day 5 post treatment was excluded from this analysis. No mortality was observed among sham-treated sires.

### Offspring survival post infection

Following the infection, sons died more slowly compared to daughters (Figure 3A; LRT, offspring sex × day post infection, χ^2^_1_ = 136.4, *p <* 0.001; Supplementary Figure S2). For further analysis, we used survival data from the day where mortality was close to 50%: day 7 post infection (DPI7) for daughters and day 9 post infection (DPI9) for sons as the measure of their pathogen resistance level. The survival of sons on DPI9 and daughters on DPI7 was moderately positively correlated across sires (Figure 3B; major axis slope = 1.416, 95% confidence interval = [0.835, 2.700]; Pearson’s correlation coefficient r = 0. 242, *p <* 0.001; see Supplementary Figure S3 for daughter-son survival relationship on each DPI). Although the daughter-son correlation is relatively low, it still suggests that genes (and possible parental effects) promoting pathogen resistance in sons also increase pathogen resistance of daughters.

**Figure 3.**
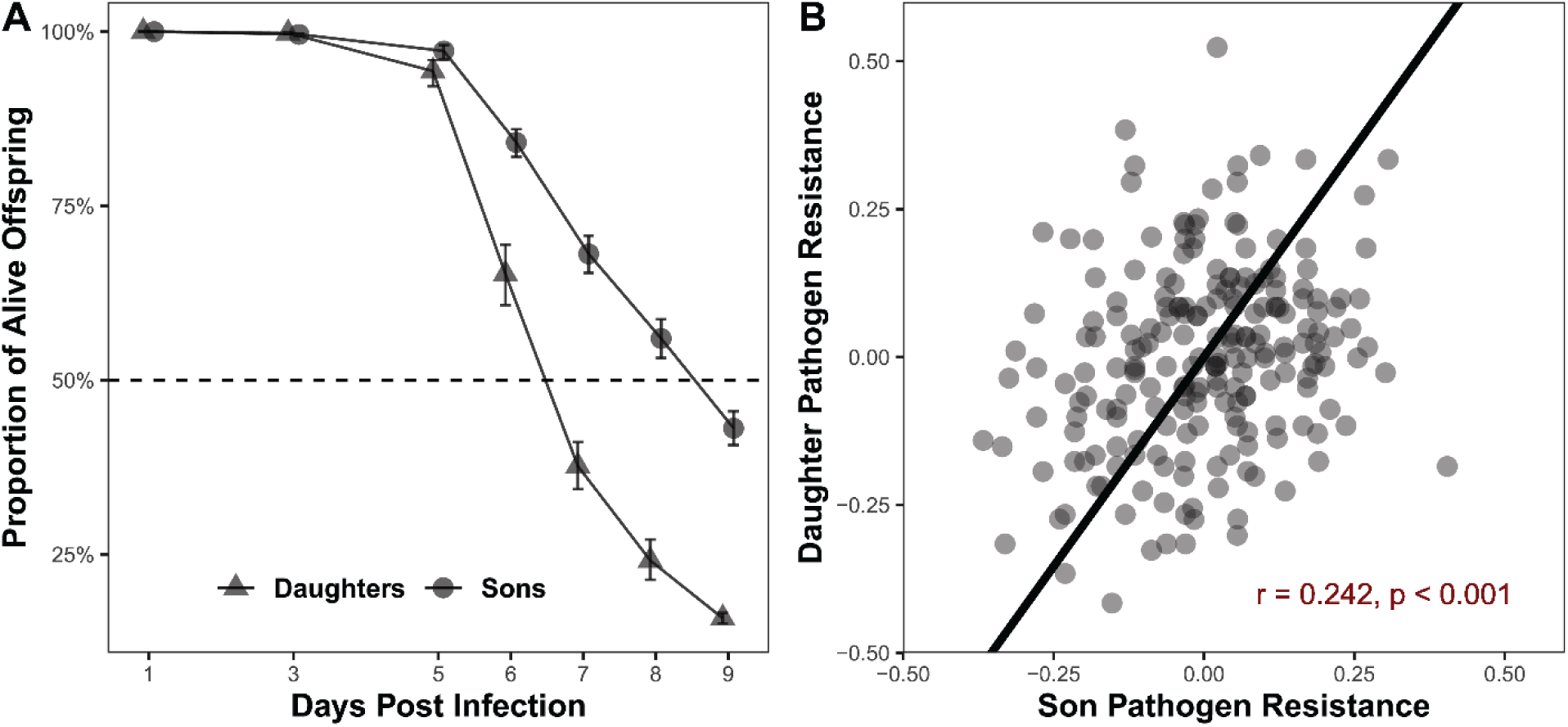
A: Survival post infection for daughters and sons (Estimated means ± SE). Estimated means are extracted from random-effect-only GLMMs (see Methods); B: Correlation between the pathogen resistance of sons and daughters of the same sire. Survival data for sons are from day 9 post infection (DPI9) and for daughters from DPI7. X- and Y-axis are zero-centered by experimental block (see Methods). Each dot is the mean of offspring coming from the same sire. Solid line displays the fitted line from the major axis regression.

### Resistant sires have higher sexual success but not more resistant offspring

Infected sires that survived beyond median post-infection survival time (i.e., those phenotypically relatively more resistant) had 13.3% higher sexual success in the mating trials than those died before the median (Figure 4A; sire pathogen resistance level: χ^2^_1_ = 4.4, *p* = 0.036). Despite weak trends, we detected no relationship between sire pathogen resistance and offspring pathogen resistance whether sons and daughters were analyzed separately (pairwise comparison, relatively resistant sire vs. relatively susceptible sire, son, *p* = 0.89, daughter, *p* = 0.22; Figure 4B, C) or together (*p*=0.48; Supplementary Figure S4).

**Figure 4.**
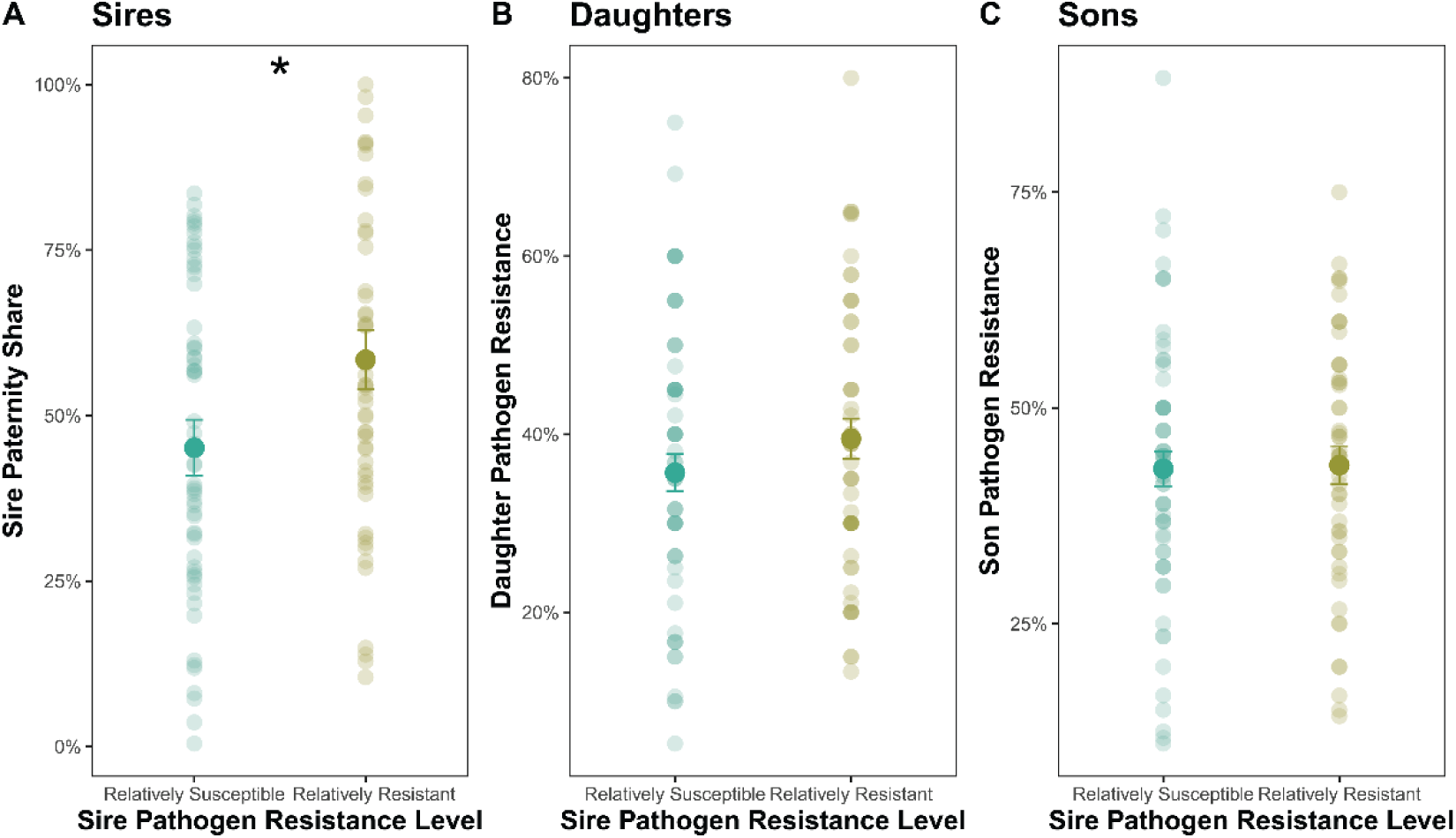
Relationship between sire’s pathogen resistance level with A: sire’s paternity share; B: daughter’s pathogen resistance and C: son’s pathogen resistance. Each dot in A represents measures from each sire and in B-C represents the survival of ∼20 offspring from each sire. Symbols show the estimated marginal means ± SE; *p ≤ 0.05.

### Sire-offspring relationship is offspring-sex-specific and depends on the pathogenic environment

The relationship between sire sexual success and offspring pathogen resistance varied depending on the sire’s treatment prior to the mating trial and offspring sex (GLMM, sire infection treatment × paternity share × offspring sex, χ^2^_1_ = 5.4, *p* =0.021; see Supplementary Table S4 for the complete model statistics). Therefore, further analyses were split by offspring sex to test the relationship between sire sexual success and pathogen resistance of sons and daughters. For sons, the most parsimonious GLMM describing the data includes sire paternity share, sire infection treatment, and their interaction as fixed factors. For daughters, it includes sire paternity share, mating scenario, and their interaction as fixed factors. Both GLMMs include experiment block as an additional fixed factor and sire identity as the random factor.

For sons, the mating scenario did not significantly affect the relationship between their pathogen resistance and sire sexual success (the best model fitting the data is a GLMM without the mating scenario), which is also confirmed by the similar sire-son relationship seen in the Long Mating scenario and the Short Mating scenario (Figure 5A). We found a significant interaction between sire’s paternity share (i.e., sexual success) and sire treatment was (Figure 5A; χ^2^_1_ = 4.2, *p* =0.041; Supplementary Table S5). For sires subject to sham treatment prior to mating trial, sires with higher paternity share had sons of lower post infection survival (paternity share, χ^2^_1_ = 6.8, *p* = 0.009; estimated slope (log-odds scale) ± SE = –0.72 ± 0.27), but when sires were infected prior to the mating trial, such negative correlation was no longer detectable (paternity share, χ^2^_1_ = 0.0, *p* = 0.99; estimated slope ± SE = –0.00 ± 0.25).

**Figure 5.**
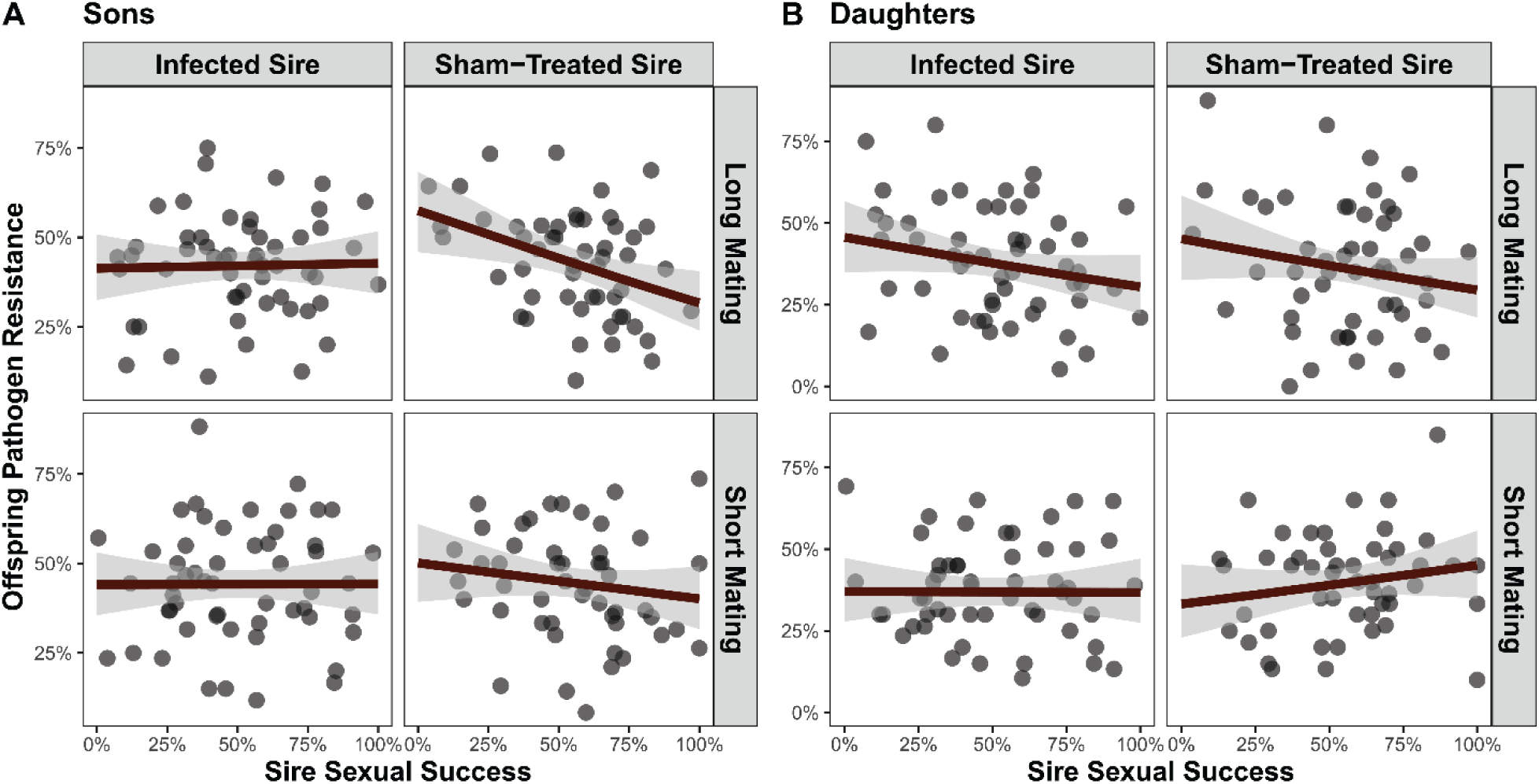
Relationship between sire sexual success and pathogen resistance of A: Sons; B: Daughters. Each dot represents the mean of ∼20 offspring from the sire. Solid lines represent the predicted values of a generalized linear mixed model (sire’s infection treatment, mating scenarios and paternity share as fixed factors; experiment block and observation identity as the random factors). Grey shadows around lines indicate 95% confidence intervals.

For daughters, sire infection treatment did not significantly affect the relationship between their survival and the sire’s paternity share (the best model fitting the data is the GLMM without sire infection treatment), but the sire-daughter relationships observed in the two mating scenarios were different (Figure 5B; paternity share × mating scenario, χ^2^_1_ = 4.7, *p* =0.030; Supplementary Table S6). When measured in the Long Mating scenario, sire’s paternity share was tended to be negatively correlated with the daughters’ pathogen resistance regardless of the sire’s infection treatment (paternity share, χ^2^_1_ = 3.7, *p* =0.056; estimated slope ± SE = -0.68 ± 0.35). In the Short Mating scenario, we found no correlation between daughters’ pathogen resistance and sire sexual success regardless of whether sires were subjected to infection prior to the mating trial (paternity share, χ^2^_1_ = 1.1, *p* = 0.30; estimated slope ± SE = 0.25 ± 0.24, *p* = 0.30).

## Discussion

In this study, we set out to test the hypothesis that the relationship between sire sexual success and offspring pathogen resistance would depend on the epidemiological context under which sexual selection takes place. Specifically, we predicted that this correlation would be positive if the sires were exposed to the pathogen prior to the action of sexual selection, but none or negative otherwise. We found that this focal relationship indeed depended on the sire pathogen exposure, but the pattern turned out to be more complicated than our relatively simple prediction. The first and most significant pattern we found is that the sire-offspring relationship was different for daughters and for sons. Furthermore, the factors retained in the most parsimonious models describing the sire-offspring relationship differed between sexes, indicating that the consequences of sexual selection on daughters and sons depend on different circumstances.

For sons, the sign of the sire-son relationship was affected by sires’ pathogen exposure. When sires were sham-treated prior to mating trials, sires that were more sexually successful had sons that were more susceptible to the fungal infection. This implies a cost of carrying resistant alleles in a pathogen-free environment, which aligns with other studies that have demonstrated the trade-offs between immunity and other fitness components, e.g., longevity (Hunter, 2011), fecundity (McKean & Lazzaro, 2011) and male’s mating success (Kawecki, 2020). However, when sires were exposed to pathogens before entering the mating trial, no correlation between sires and sons was detected. This sire-son relationship which varies with epidemiological context , partially supports the “context-dependent” hypothesis (Westneat & Birkhead, 1998) and is consistent with findings from Joye and Kawecki (2019). It also illustrates that pathogen exposure modulates the effects of resistant genetic variants on sexual success, adding to the existing empirical evidence showing genotype by environment interactions in sexual selection (Danielson-Francois et al., 2006; Engqvist, 2008; Ingleby et al., 2010; Lewis et al., 2012; Ingleby et al., 2013). We found no “good genes” for sons with respect to pathogen resistance as we did not find any significantly positive sire-son relationship in either mating scenario, whether or not when pathogen was present.

For daughters, we did not find any evidence for “good genes” either. In the Long Mating scenario, regardless of sire infection treatment, daughters of more sexually successful sires tended to be somewhat more susceptible. We speculate that the reason behind this somewhat negative sire-daughter relationship detected when sires were not exposed to the pathogen instead of the no correlation seen for sons is likely due to sexually antagonistic selection. It could be that some of the genes favored by sexual selection on sires have antagonistic effects on daughters or at best, sexually selected genes in sons may not be as beneficial to daughters (sex-specific effects), as has been reported in Guncay et al. (2017) and Joye and Kawecki (2019), respectively. Positive covariance between the pathogen resistance of sons and daughters indicates that most of genetic variants affect resistance of males and females similarly. However, some of the sons and daughters were full sibs, so dominance genetic variance also contributes to the son-daughter correlation but not the sire-offspring correlation. Furthermore, a moderate positive correlation is not incompatible with the existence of some variants that have sex-specific or sex-antagonistic effects, and it could be that these variants are linked to male sexual success.

The fact that the mating scenario (i.e., length of interactions in the mating trial) had a stronger impact on the sire-daughter relationship than on the sire-son relationship further supports our speculation on the sexually antagonistic selection. In the Long Mating scenario, the captured sire sexual success reflects sexual interactions with (re)mated females, whereas competition for virgin females is more common in the Short Mating scenario. Mated females are less willing to mate and more discerning about male traits (Jennions & Petrie, 2000; Kokko & Mappes, 2005; Kohlmeier et al., 2021), thus resulting in a potentially stronger preference for male-benefit traits in the Long Mating scenario and potentially a stronger antagonistic gene effects on daughters.

The “good genes” hypothesis and its more specific variants like the Hamilton-Zuk hypothesis have been initially proposed to explain female preference. However, an extended duration of sexual interactions and at high-density lab conditions may have left females limited opportunities to escape constant attention/harassment from males and to exercise choice freely. That is, in our study, females might have still preferred to mate with males that carry genetic variants for resistance, but may not have had the freedom to exercise this preference – sexual conflict over mating may negate or even overwrite the genetic benefits from female preference (Snow et al., 2019; Yun et al., 2021; Flintham et al., 2023). A briefer sexual contact like in the Short Mating scenario may to some extent relieve such stress on the females or a more structured environment allowing or lower density, allowing females to escape harassment (Sharp & Agrawal, 2008; Yun et al., 2017; Talagala et al., 2024).

Our experiment design ensured that the most probable explanation for the complex but significant pattern of the relationship between sire paternity share and offspring post infection survival is that there is additive genetic covariance. Nonetheless, despite a trend, we did not detect a positive relationship between sire and offspring pathogen resistance, suggesting that the additive genetic variance is rather low. Although sires of higher sexual success also had higher resistance against *Metarhizium*, such positive correlation on the phenotypic level does not reflect an additive genetic correlation. Males may be more sexually successful and more resistant to pathogens because they carry a higher level of heterozygosity, which is a non-additive genetic component (Brown, 1997; Kempenaers, 2007). Another possible explanation is that sire pathogen resistance was measured within a sexual selection context, which may strengthen the immunity-reproduction trade-off (Kawecki, 2020). In contrast, for practical reasons offspring pathogen resistance was measured in single sex groups; excluding the potential confounding effect of reproductive effort on survival.

The “good genes” hypothesis of sexual selection is often thought to reinforce the alignment of sexual selection and natural selection and to increase the population mean fitness in the long run. Nonetheless, we found no support for “good genes” sexual selection promoting pathogen resistance under any circumstances in our study, echoing findings from Sharda et al. (2022) that sexual selection did not promote resistant genes in the population after 12 generations of experimental evolution. In nature, males get infected and continue to interact with the other males and females during the course of infection, which is similar to what has been shown in our Long Mating scenario. The context- and sex-dependent sire-offspring relationship we detected here demonstrates that the cost-benefit balance of carrying pathogen resistance was mediated by the context where sexual selection happens and how sex-specific or sexually antagonistic selection may break the alignment of the two forms of selections on promoting resistant genes (Rice & Chippindale, 2001; Pischedda & Chippindale, 2006; Foerster et al., 2007; Hollis & Houle, 2011). In particular, our results reveal the complex interplay between sexual selection and natural selection, and highlight the need to test the “good genes” hypothesis and the different forms of genotype by environment interactions on a larger temporal and spatial scale (Cornwallis & Uller, 2010).

## Data accessibility

All data and R scripts are available on Zenodo (https://doi.org/10.5281/zenodo.13355838).

## Author contributions

A.L. and T.J.K. conceptualized the study; A.L. collected the data; A.L. and T.J.K. wrote the manuscript.

## Competing interests

We declare no competing interests.

## Funding

This work is supported by the Swiss National Science Foundation research grant 310030_184791 to TJK.

## Acknowledgements

We thank multiple student research assistants in the Kawecki lab for preparing hundreds of bottles/ vials of fly food for our experiment. We would like to thank Dr. Luc Bussière for commenting on the early draft of this manuscript.

## Notes

### Competing Interest Statement

The authors have declared no competing interest.

