## Supplementary for "Context- and sex-dependent links between sire sexual success and offspring pathogen resistance"

### Table of Contents


Supplementary Figure S 1 Survival curves of infected sires following infection under the two mating scenarios. Arrows indicate splitting time points for sire pathogen resistance levels: relatively susceptible (sires dead before or on the splitting time point) and relatively resistance (sires alive after the splitting time point). The splitting time point is different for “Long Mating” and “Short Mating” scenario. ............ 3

Supplementary Figure S 2 Survival of daughters and sons post infection (means ± SE). Different experiment blocks are indicated by different line types. .................................................................. 4

Supplementary Figure S 3 Relationship between son survival and daughter survival on each day post infection. Each dot represents the offspring from the same sire. x axis and y axis indicate the difference between the observed survival% to the mean offspring survival% of the corresponding block. Solid lines are the fitted lines from linear regression. Pearson’s correlation coefficient (R) and the corresponding significance level are also shown on the figure. .................................................................. 5

Supplementary Figure S 4 Relationship between sire’s pathogen resistance level with combined offspring pathogen resistance. Each dot in represents the survival of ~40 offspring from each sire. Error bars show the estimated marginal means ± SE. .................................................................. 6

Supplementary Table S 1 Sample size (number of sires) of the main experiment .................................... 7

Supplementary Table S 2 Likelihood ratio tests of fixed factors on paternity share. The generalized linear mixed model includes paternity share (i.e., number of embryos sired by focal male versus that sired by standard competitors) as the response variable, sire infection treatment, days post treatment and their interaction, and experiment block (N = 3) as fixed factors and focal sire identity as the random factor. Significance terms ($p \leq 0.05$) are indicated in bold. .................................................................. 8

Supplementary Table S 3 Estimated slopes of the relationship between days post treatment and paternity share for infected and sham-treated sires, and the statistical summary of the pairwise contrast (supplementary to Supplementary Table S2). Significance terms ($p \leq 0.05$) are indicated in bold, LCL and UCL refer to lower and upper 95% confidence limits. .................................................................. 8

Supplementary Table S 4 Effects of the fixed factors on offspring survival based on the Likelihood Ratio Test (**Whole dataset**). The generalized linear mixed model includes offspring survival as the response variable, sire infection treatment, paternity share, mating scenario, offspring sex and their interactions and experiment block as fixed factors, and focal sire identity and observation identity as the random factors. Significance terms ($p \leq 0.05$) are indicated in bold. .................................................................. 9

Supplementary Table S 5 Effects of the fixed factors based on the likelihood ratio tests (**Sons only dataset**). The best GLMM fitting to the data (with the lowest AIC) includes offspring survival as response variable, sire infection treatment, paternity share and their interaction, and experiment block as fixed factors and sire identity as an observational level random factor to correct for overdispersion. Significance terms ($p < 0.05$) are indicated in bold. .................................................................. 10

Supplementary Table S 6 Effects of the fixed factors based on the likelihood ratio tests (**Daughters only dataset**). The best GLMM fitting to the data (with the lowest AIC) includes offspring survival as response variable, mating scenario, paternity share and their interaction, and experiment block as fixed factors and sire identity as an observational level random factor to correct for overdispersion. Significance terms ($p < 0.05$) are indicated in bold. .................................................................. 10

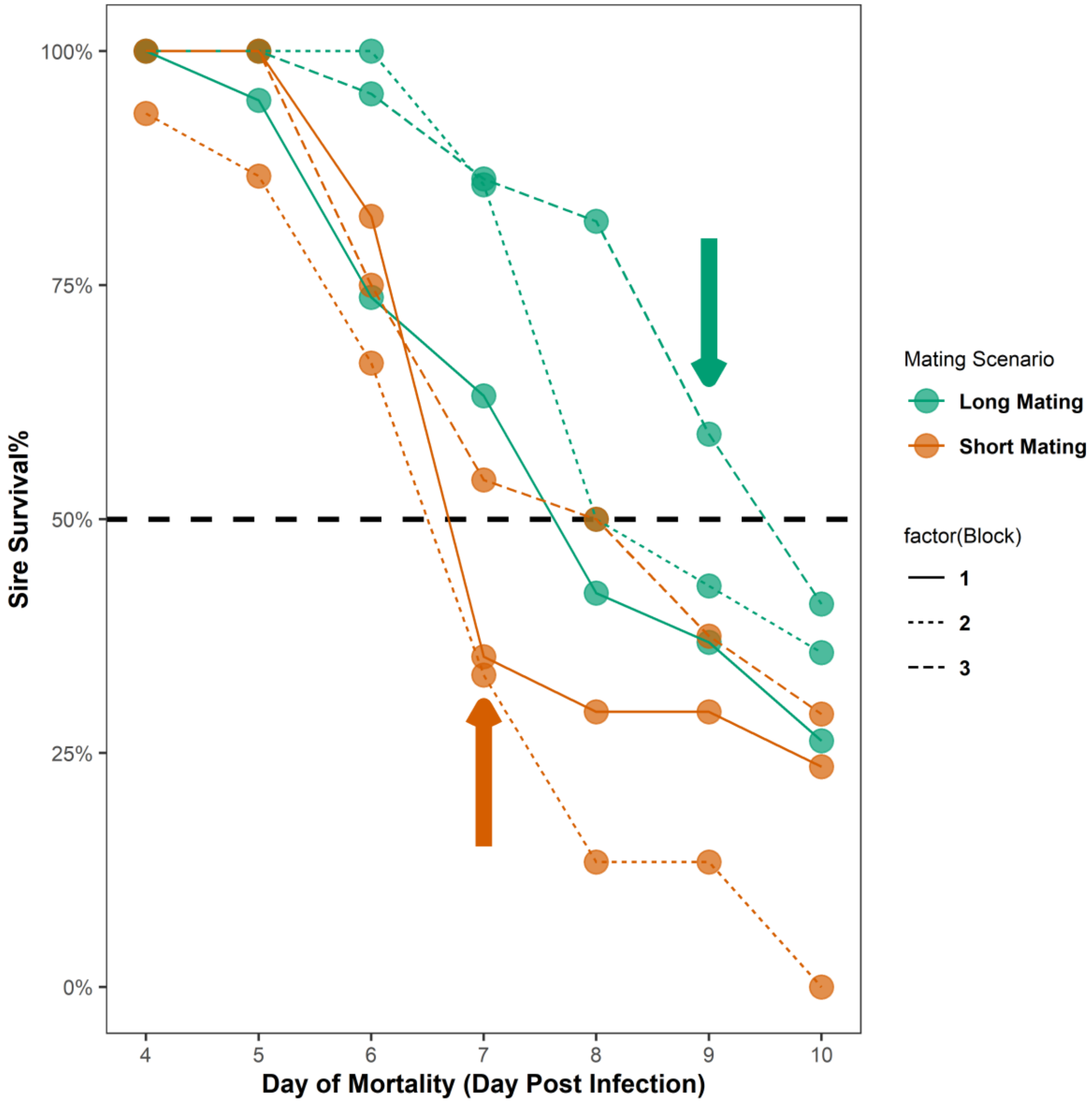


*Supplementary Figure S 1 Survival curves of infected sires following infection under the two mating scenarios. Arrows indicate splitting time points for sire pathogen resistance levels: relatively susceptible (sires dead before or on the splitting time point) and relatively resistance (sires alive after the splitting time point). The splitting time point is different for "Long Mating" and "Short Mating" scenario.*

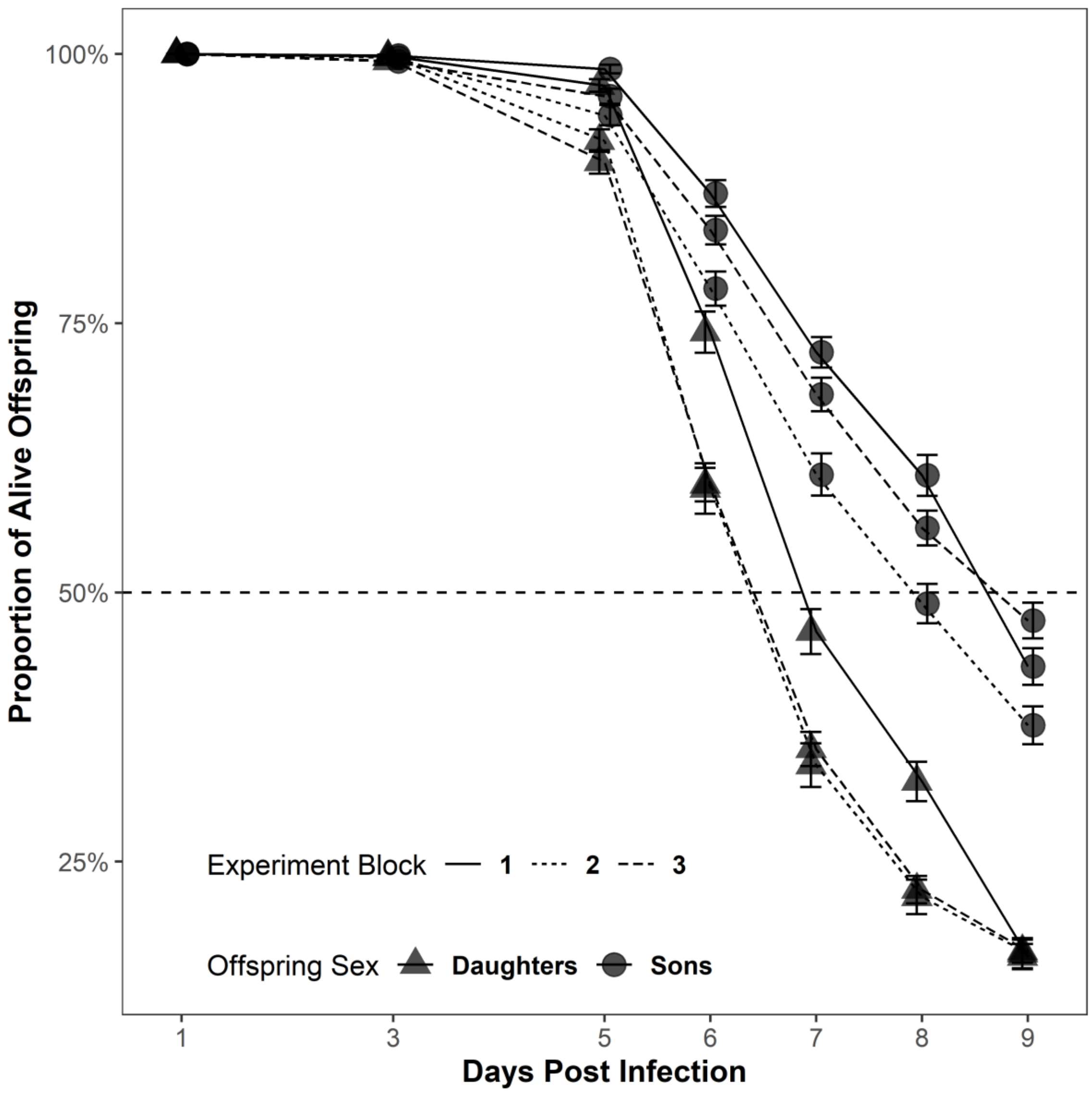


*Supplementary Figure S 2 Survival of daughters and sons post infection (means ± SE). Different experiment blocks are indicated by different line types.*

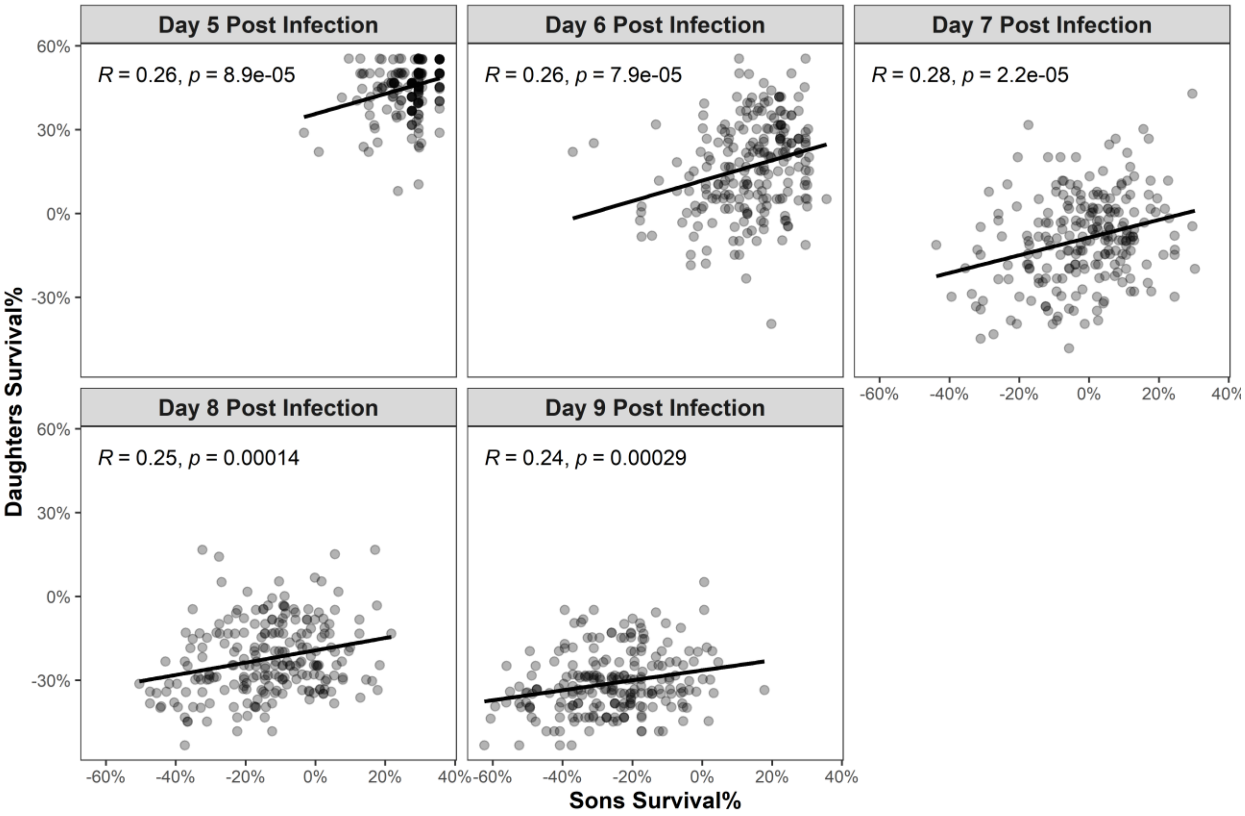


*Supplementary Figure S 3 Relationship between son survival and daughter survival on each day post infection. Each dot represents the offspring from the same sire. x axis and y axis indicate the difference between the observed survival% to the mean offspring survival% of the corresponding block. Solid lines are the fitted lines from linear regression. Pearson's correlation coefficient (R) and the corresponding significance level are also shown on the figure.*

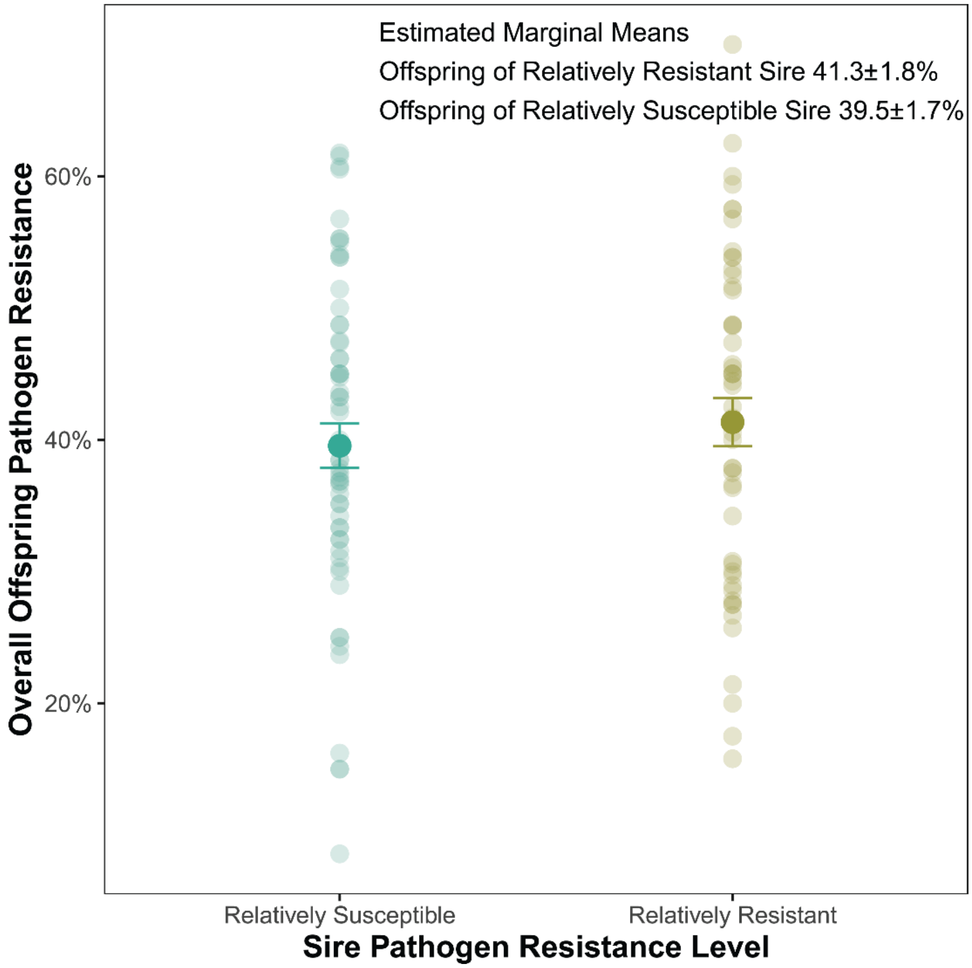


*Supplementary Figure S 4 Relationship between sire's pathogen resistance level with combined offspring pathogen resistance. Each dot in represents the survival of ~40 offspring from each sire. Error bars show the estimated marginal means ± SE.*

*Supplementary Table S 1 Sample size (number of sires) of the main experiment*

| Experiment Block | Block 1 | | Block 2 | | Block 3 | |
|---|---|---|---|---|---|---|
| Mating Scenario | Long Mating | Short Mating | Long Mating | Short Mating | Long Mating | Short Mating |
| Infected | 18 | 18 | 15 | 14 | 22 | 24 |
| Sham-treated | 15 | 16 | 13 | 14 | 24 | 19 |

*Supplementary Table S 2 Likelihood ratio tests of fixed factors on paternity share. The generalized linear mixed model includes paternity share (i.e., number of embryos sired by focal male versus that sired by standard competitors) as the response variable, sire infection treatment, days post treatment and their interaction, and experiment block (N = 3) as fixed factors and focal sire identity as the random factor. Significance terms (p ≤ 0.05) are indicated in bold.*

| Factor | $\chi 2$ | d.f. | *p* |
|---|---|---|---|
| Long Mating (DPT4-6) | | | |
| Sire Infection Treatment | 0.9 | 1 | 0.343 |
| **Days Post Treatment** | **318.5** | **1** | **< 0.001** |
| **Experiment Block** | **9.5** | **2** | **0.009** |
| **Treatment: Days Post Treatment** | **43.5** | **1** | **< 0.001** |
| Short Mating (DPT4-6) | | | |
| Sire Infection Treatment | 3.6 | 1 | 0.058 |
| **Days Post Treatment** | **330.8** | **1** | **< 0.001** |
| Experiment Block | 2.2 | 2 | 0.331 |
| **Treatment: Days Post Treatment** | **12.3** | **1** | **< 0.001** |

*Supplementary Table S 3 Estimated slopes of the relationship between days post treatment and paternity share for infected and sham-treated sires, and the statistical summary of the pairwise contrast (supplementary to Supplementary Table S2). Significance terms (p ≤ 0.05) are indicated in bold, LCL and UCL refer to lower and upper 95% confidence limits.*

| Long Mating (DPT4-6) | | | | |
|---|---|---|---|---|
| Sire Infection Treatment | Slope | SE | LCL | UCL |
| Infected | -0.266 | 0.016 | -0.296 | -0.235 |
| Sham | -0.122 | 0.015 | -0.152 | -0.093 |
| Pairwise Contrast | | | | |
| contrast | estimate | SE | *z* | *p* |
| **Infected-Sham** | **-0.144** | **0.022** | **-6.595** | **< 0.001** |

| Short Mating (DPT4-6) | | | | |
|---|---|---|---|---|
| Treatment | Slope | SE | LCL | UCL |
| Infected | -0.333 | 0.023 | -0.377 | -0.289 |
| Sham | -0.225 | 0.021 | -0.267 | -0.184 |
| Pairwise Contrast | | | | |
| contrast | estimate | SE | *z* | *p* |
| **Infected-Sham** | **-0.108** | **0.031** | **-3.502** | **< 0.001** |

*Supplementary Table S 4 Effects of the fixed factors on offspring survival based on the Likelihood Ratio Test (**Whole dataset**). The generalized linear mixed model includes offspring survival as the response variable, sire infection treatment, paternity share, mating scenario, offspring sex and their interactions and experiment block as fixed factors, and focal sire identity and observation identity as the random factors. Significance terms ($p \leq 0.05$) are indicated in bold.*

**FULL MODEL (Whole dataset)**

| Factor | $\chi 2$ | d.f. | *p* |
|---|---|---|---|
| Sire Infection Treatment | 0.3 | 1 | 0.614 |
| Paternity Share | 3.2 | 1 | 0.075 |
| **Offspring Sex** | **15.4** | **1** | **< 0.001** |
| Mating Scenario | 0.7 | 1 | 0.392 |
| **Experiment Block** | **20.2** | **2** | **< 0.001** |
| Sire Infection Treatment × Paternity Share | 0.7 | 1 | 0.394 |
| Sire Infection Treatment × Offspring Sex | 0.0 | 1 | 0.969 |
| Paternity Share × Offspring Sex | 0.1 | 1 | 0.812 |
| Sire Infection Treatment × Mating Scenario | 0.1 | 1 | 0.756 |
| Paternity Share × Mating Scenario | 3.7 | 1 | 0.054 |
| Offspring Sex × Mating Scenario | 0.0 | 1 | 0.919 |
| **Sire Infection Treatment × Paternity Share × Offspring Sex** | **5.4** | **1** | **0.021** |
| Sire Infection Treatment × Paternity Share × Mating Scenario | 0.9 | 1 | 0.343 |
| Sire Infection Treatment × Offspring Sex × Mating Scenario | 0.8 | 1 | 0.367 |
| Paternity Share × Offspring Sex × Mating Scenario | 2.0 | 1 | 0.153 |
| Sire Infection Treatment × Paternity Share × Offspring Sex × Mating Scenario | 0.2 | 1 | 0.685 |

*Supplementary Table S 5 Effects of the fixed factors based on the likelihood ratio tests (**Sons only dataset**). The best GLMM fitting to the data (with the lowest AIC) includes offspring survival as response variable, sire infection treatment, paternity share and their interaction, and experiment block as fixed factors and sire identity as an observational level random factor to correct for overdispersion. Significance terms ($p < 0.05$) are indicated in bold.*

| Factor | χ2 | d.f. | *p* |
|---|---|---|---|
| Sire Infection Treatment | 0.2 | 1 | 0.666 |
| Paternity Share | 3.5 | 1 | 0.061 |
| **Experiment Block** | **20.0** | **2** | **< 0.001** |
| **Sire Infection Treatment × Paternity Share** | **4.2** | **1** | **0.041** |

*Supplementary Table S 6 Effects of the fixed factors based on the likelihood ratio tests (**Daughters only dataset**). The best GLMM fitting to the data (with the lowest AIC) includes offspring survival as response variable, mating scenario, paternity share and their interaction, and experiment block as fixed factors and sire identity as an observational level random factor to correct for overdispersion. Significance terms ($p < 0.05$) are indicated in bold.*

| Factor | χ2 | d.f. | *p* |
|---|---|---|---|
| Paternity Share | 1.0 | 1 | 0.317 |
| Mating Scenario | 0.3 | 1 | 0.569 |
| **Experiment Block** | **24.1** | **2** | **< 0.001** |
| **Paternity Share × Mating Scenario** | **4.7** | **1** | **0.030** |
